# Genome Instability Precedes Viral Integration in Human Papilloma Virus Transformed Tonsillar Keratinocytes

**DOI:** 10.1101/2024.08.14.607944

**Authors:** Kimberly Chan, Christopher Tseng, Emily Milarachi, David Goldrich, Lisa Schneper, Kathryn Sheldon, Cesar Aliaga, Samina Alam, Sreejata Chatterjee, Karam El-Bayoumy, Craig Meyers, David Goldenberg, James R. Broach

**Author notes:** Corresponding Author: James R. Broach, Department of Biochemistry, Penn State College of Medicine, Hershey PA 17033. **Competing Interest Statement:** The authors declare no potential conflicts of interest.

## Abstract

Approximately 70% of oropharyngeal squamous carcinomas (OPSCC) are associated with human papillomavirus (HPV). While HPV-positive (HPV+) OPSCC responds better to standard therapies and patients with HPV+ tumor generally have better outcomes than those with HPV-negative (HPV-) tumors, a subset of HPV+ patients do have poor outcomes. Our previous work suggested that tumors with integrated virus exhibit significantly greater genome wide genomic instability than those with only episomal viral genomes and patients with HPV+ OPSCC with episomal viral genomes had better outcomes. To explore the causal relation between viral integration and genomic instability, we have examined the time course of viral integration and genetic instability in tonsillar keratinocytes transformed with HPV16. HPV-infected human tonsil keratinocyte cell lines were continuously passaged and every fifth passage some cells were retained for genomic analysis. Whole genome sequencing and optical genomic mapping confirmed that virus integrated in five of six cell lines while remaining episomal in the sixth. In all lines genome instability occurred during early passages, but essentially ceased following viral integration but continued to occur later passages in the episomal line. To test tumorigenicity of the cell lines, cells were injected subcutaneously into the flanks of nude mice. A cell line with the integrated virus induced tumors following injection in the nude mouse while that with the episomal virus did not. We conclude that genomic instability is not the result of viral integration but likely promotes integration. Moreover, those transformants with episomal virus appear to be less tumorigenic than those with integrated virus.

## Introduction

Head and Neck Squamous Cell Carcinoma (HNSCC), an aggressive malignancy with high morbidity and mortality, is the seventh most common cancer worldwide (1–5). While the classic risk factors have been tobacco and alcohol, human papillomavirus (HPV) has emerged as the major risk factor for some of these cancers, specifically for oropharyngeal squamous cell carcinoma (OPSCC). The fraction of head and neck cancers diagnosed as HPV-positive (HPV+) oropharyngeal cancers in the United States rose from 16.3% in the 1980s to more than 72.7% in the 2000s. Because of this increased prevalence of HPV+ cases, OPSCC is one of the few cancers with rapidly increasing incidence in recent years, predominantly among younger people, mostly male, and likely as a result of oral-sex exposure (6–11). Accordingly, improved methods for diagnosis, prognosis and treatment will be needed for the foreseeable future to address this epidemic of HPV-positive OPSCC.

Following initial infection, HPV persists in the nucleus of its host cell as an extrachromosomal episome, but can subsequently integrate into the host genome (12,13). The reported proportion of HPV-positive tumors in which the virus integrates into the genome varies by cohort and cancer type, with approximately 71% in virus-positive head and neck cancer and 83% in cervical cancer (14,15). These integration events occur essentially randomly throughout both the human and viral genomes. Most often only a fragment of the viral genome is retained following integration, spanning E6, E7 and a random amount of the adjacent viral genome but lacking an intact E2 (15,16). The retention of E6 and E7 and the loss of E2 is likely the consequence of selection during tumorigenesis, as the elimination of E2 results in increased expression of the E6/E7 viral oncogenes, which drives tumorigenesis. The E6 protein of the oncogenic strains of HPV inactivates the p53 pathway by promoting degradation of p53, resulting in abrogation of genome integrity surveillance (17,18). In addition, the second HPV oncoprotein, E7, binds to and inactivates the cell cycle inhibitor, Rb, leading to unrestricted cell cycle progression. Accordingly, tumorigenesis results predominantly from loss of cell cycle regulation elicited by E7 and abrogation of DNA damage checkpoint control caused by E6.

The HPV status of OPSCC tumors influences patients’ outcomes. In general, overall survival and disease-free survival are more favorable for patients with HPV+ OPSCC, who tend to have better responses to surgery and radiotherapy (19,20). However, several studies have noted outcome differences in subcategories of HPV+ HNSCCs. For instance, Jung et al (21) noted that patients whose tumors exhibit high E6/E7 and p16 expression have better outcomes than other HPV+ cases. Similarly, Mooren et al. (22) observed better outcomes in HPV+ tonsillar squamous cell carcinoma versus HPV-negative but that chromosome instability, even in the HPV+ cases, correlates with unfavorable prognosis. In fact, around 20% of HPV+ OPSCC patients have poor prognoses and the defining molecular characteristics of this subset of patients have not yet been clearly identified (17). One possibility as suggested by a recent report (23) is the loss of HPV expression in subsets of cells within an HPV+ tumor, resulting in resistance to treatment and potentially worse progression-free survival. As an alternative, or potentially related hypothesis, we have proposed that one contributing factor may be the integration state of the virus (24).

We recently applied optical genome mapping (OGM) in conjunction with whole genome sequencing (WGS) to a number of OPSCC patient samples and showed that HPV integrated into the genome in two-thirds of the cases but remained episomal in the other third (24). Strikingly, in those tumors with integrated HPV the entire human genome exhibited extensive instability with multiple translocations, inversions, insertions and deletions not just at the site of integration but throughout the genome. Those cases that were HPV+ but not integrated showed little to no genome instability. This apparent correlation between genome-wide structural rearrangements and viral integration raises a question of causality: does integration induce genome instability or does genome instability lead to viral integration. The results presented here are consistent with the conclusion that genome instability precedes viral integration rather than occurring as a result of integration and that integration promotes more aggressive tumorigenesis.

## Materials and Methods

### Cell Lines and Culture Conditions

Human tonsillar keratinocytes were infected with HPV16 virus and cultured with J2 3T3 feeder cells for up to 100 passages (Figure 1) (25). Each passage represents approximately two cell doublings. At every fifth passage, aliquots of cells were frozen and some grown into organotypic (raft) cultures and assessed by hematoxylin and eosin (H&E) staining and by immunohistochemical staining for changes in morphology and gene expression associated with cancer progression (25). The E2 and E6 levels were measured by qPCR to the respective coding regions in total genomic DNA isolated from cells.

**Figure 1.**
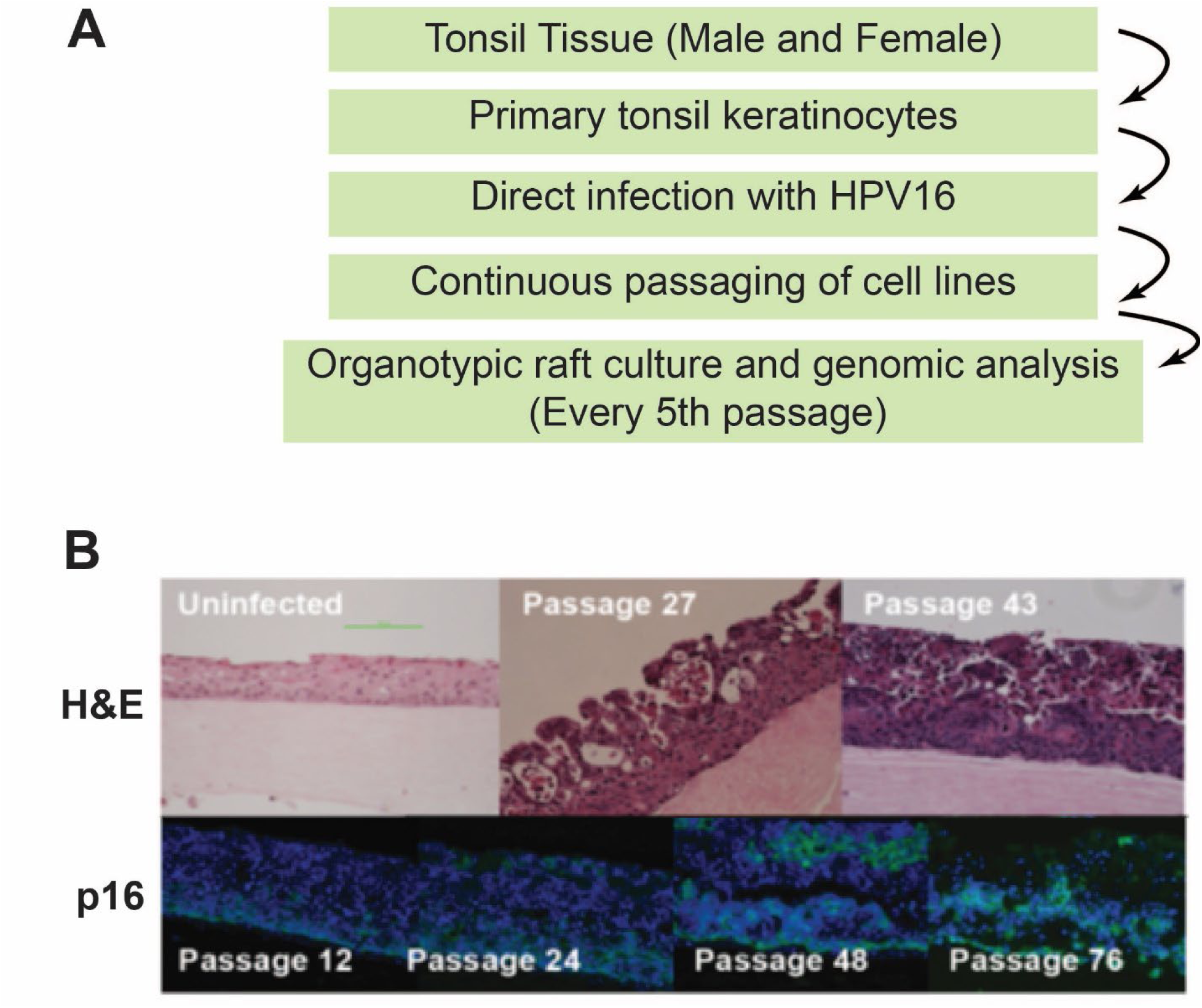
Transformation of primary tonsillar keratinocytes by HPV16. A. Scheme for generating and characterizing HPV-transformed primary tonsillar keratinocytes. B. Human tonsil keratinocytes infected with HPV16 were propagated as outlined in A. At the indicated passages a subset of cells was grown into organotypic raft cultures and either stained with hematoxylin and eosin (H&E) or immunostained for p16.

### Optical Genome Mapping (OGM)

We performed OGM on fresh cells from each cell line at six different time points during the first 50-60 passages. We extracted Ultra high molecular weight DNA from fresh cells (∼ 1.5x10^6^ as determined on an Eve Cell Counter) as recommended (CG-00037-Ionic-G2-Tissue-to-DNA-Kit-Protocol.pdf) and direct labeled it with the DLS-G2 Labeling Kit as described (bionano.com/wp-content/uploads/2024/02/CG-30553-1_Bionano-Prep-DLS-G2-Protocol.pdf). The samples were run on Bionano G2.3 Saphyr chips. The data was analyzed with the De Novo assembly using Bionano Access version 1.7. Structural variants included insertions, deletions, inversions and duplications >500 bp as well as translocations. Copy number variants were defined by gain or loss of chromosomal segments > 30 Mb. A structural variant or copy number variant that appeared in any of the passages was added to the number of accumulated SVs if the variant had an allele frequency greater than 0.1 and recurred in later passages.

### Whole Genome Sequencing

Ultrahigh molecular weight DNA previously isolated using the Bionano protocol underwent buffer exchange using the Zymo Research Genomic DNA Clean and Concentrate kit (cat no. D4033). The libraries were prepared using Illumina DNA PCR-free Prep Kit, Tagmentation according to manufacturing instructions. Unique dual-indexed adapters were used to reduce index hopping on the NovaSeq platform. Samples were run on an S1 flowcell of the Illumina NovaSeq 6000 Sequencer with 150 bp paired end reads.

### Anchored PCR

For this modification of the protocol developed by Dawes et al (26), ultrahigh molecular weight DNA (1 μg) previously isolated using the Bionano protocol was sheared to 10-20 kb using the Covaris® g-TUBE (https://www.covaris.com/wp/wp-content/uploads/2020/05/pn_010154.pdf) and then re-isolated using sparQ PureMag beads (sparQ beads). DNA libraries were prepared with the KAPA HyperPrep Kit using custom 15 μM half-adapters and then isolated using sparQ beads. PCR amplification using the Phusion® Hot Start Flex DNA Polymerase was performed in either 25 μL reaction volume for 20-30 ng of library DNA or 50 μL for 40-50 ng of library DNA (https://www.neb.com/en-us/protocols/2012/09/06/protocol-for-phusion-hot-start-flex-dna-polymerase-m0535). Each sample was performed in triplicate with indexed barcoded primers: one with the E6 primer, one with the reverse E6 primer (E6v2), and one no template control (NTC). The reactions incubated in the thermocycler for 16 cycles.

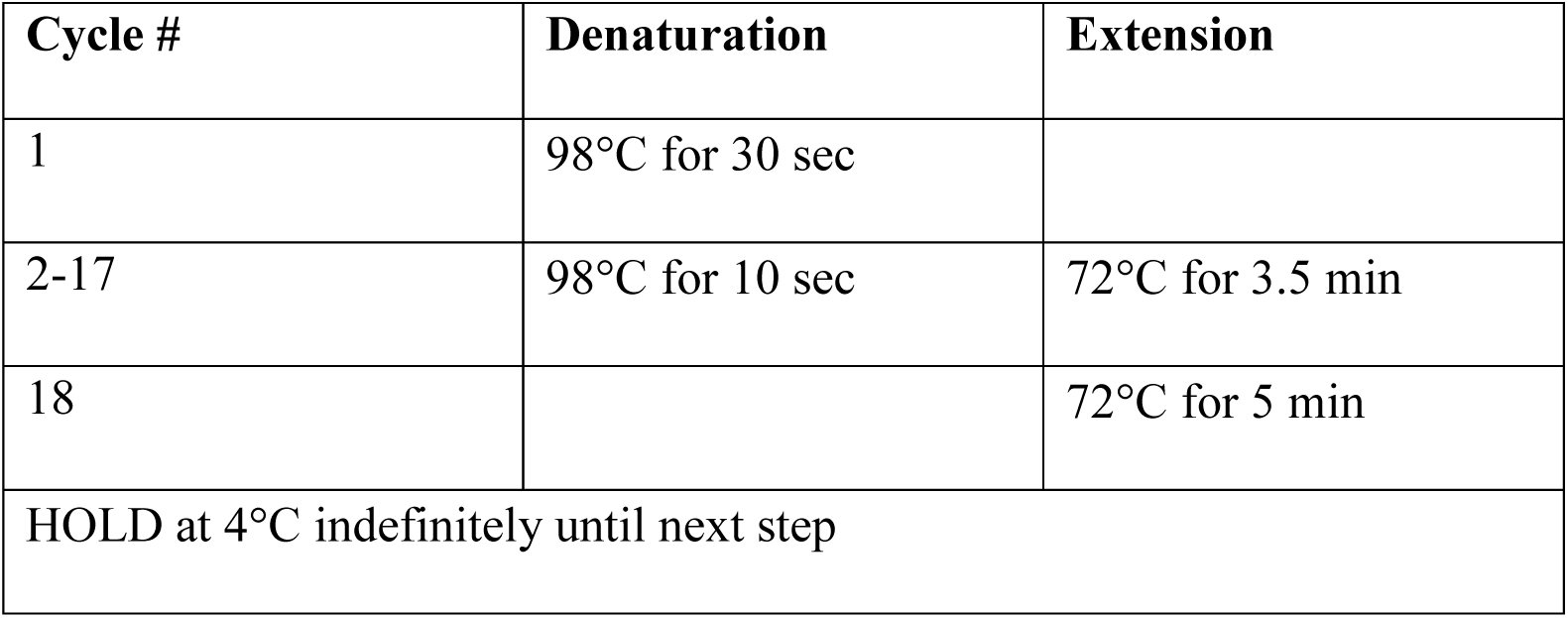

| Cycle # | Denaturation | Extension |
| --- | --- | --- |
| 1 | 98°C for 30 sec |  |
| 2-17 | 98°C for 10 sec | 72°C for 3.5 min |
| 18 |  | 72°C for 5 min |
| HOLD at 4°C indefinitely until next step |  |  |

The reactions were cleaned up using sparQ beads and then were quantified using HS Qubit. The DNA 12000 kit was used to analyze the samples on the Bioanalyzer to ensure the amplified DNA were of adequate size. The samples were pooled and HS Qubit was performed to obtain an approximate final concentration before sequencing. Samples were sequenced on PacBio® Sequel IIe System, after confirming that the amplified DNA fragments were 10 Kbp or greater by analysis of an aliquot on an Agilent TapeStation 4150.

### Assessment of tumor growth in nude mice

Seven weeks old Athymic nude Crl:NU(NCr)-Foxn1nu immunodeficient outbred female mice were obtained from Charles River Laboratories. After a week of acclimation, mice were injected in the right and left flank with 1x10^7^ cells suspended in 200 μl PBS. Prior to injection, cells were cultured for three days in E medium in the presence of mitomycin C-treated J2 3T3 feeder cells (27) with or without 10 nM dibenzo[def,p]chrysene-11,12-dihydrodiol 13,14-epoxide (synthesized by the Penn State Cancer Institute Organic Synthesis Core as described (28)). At 70% confluence, cells were trypsinized, pelleted, washed with PBS and pelleted again and resuspended in PBS. Treated cells were injected on the left flank and untreated on the right of five separate mice for each strain. Two dimensional measurements of the tumors were begun seven days after inoculation and repeated twice a week for four weeks. Volumes were calculated as Length x (Width/2)^2^.

## Results

### HPV often integrates during propagation of HPV-transformed primary tonsillar keratinocytes

In order to examine the events occurring around the time of viral integration during HPV-induced tumorigenesis, we examined transformants obtained by HPV infection of primary tonsillar keratinocytes as previously described (Figure 1A) (25). Briefly, six HPV16 infected human tonsil keratinocyte cell lines, three from male donors and three from female, were continuously passaged and every fifth passage some cells were retained for subsequent use. Additionally, at various intervals some cells were grown in organotypic raft culture to assess changes in morphology and gene expression patterns normally associated with cancer progression. Expression of p16, a prognostic marker of HPV16 oropharyngeal cancer (29,30), increased in this progression model (Figure 1B). These observations confirm that the transformed cells obtained by HPV infection exhibited features characteristic of OPSCC. In addition, we monitored the state of the HPV genome to assess its integration status. The ratio of E2 to E6 DNA provided an indirect indicator of integration, since as noted above E2 is almost always disrupted during integration while E6 and E7 must remain intact to maintain the transformed state. By this measure, HPV became integrated during passage of some of the cell lines but remained episomal in others (Figure 2).

**Figure 2.**
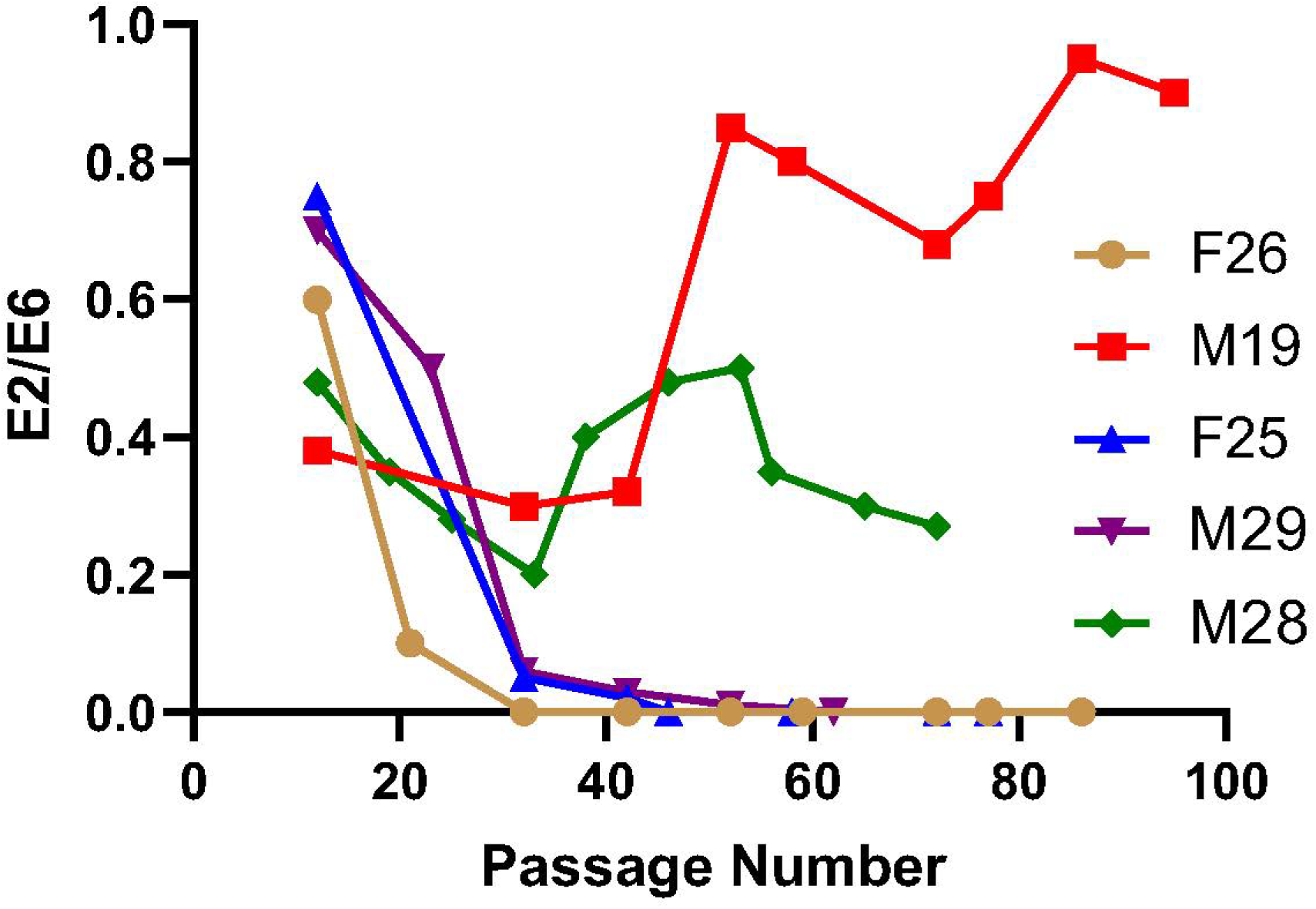
Assessment of HPV integration status by E2/E6 ratios. Shown are HPV E2/E6 ratios as a function of passage number during propagation of five independent human tonsillar keratinocyte cell lines following infection with HPV16. E2 and E6 levels were determined by qPCR.

To confirm and characterize the integration events, we performed OGM on cells taken at approximately every tenth passage of six separate cell lines, the five noted in Figure 2 and one additional line. We also performed WGS on cells obtained from approximately the forty-fifth passage of each cell line, since data in Figure 2 indicated that, for those lines in which HPV apparently integrated, integration occurred by that passage. To conclude that the virus had integrated we required that WGS returned a significant number of hybrid reads in which viral sequences abutted human genomic sequences as well as a number of split reads in which one of the paired end reads mapped to the virus and the other paired end read mapped to the human genome at a site consistent with the hybrid reads. We further required that the OGM data for cells from the same passage showed an insertion at the site predicted by the WGS reads. By these criteria the virus had integrated into the host genome in five of the six cell lines. Moreover, consistent with previous results from tumor tissue, viral integration resulted in loss of E2 and retention of E6 and E7 (Figure 3). In four of the five cell lines, this occurred through deletion a significant fraction of the viral genome eliminating E2 but retaining E6 and E7. In one interesting case, the entire viral genome was retained with a duplication of the E2 coding region but with truncation of opposite ends of the duplicated gene, rendering both copies of E2 likely inactive. This likely derived from a viral genome dimer and accounts for the discrepancy in integration status of the virus as determined by E2/E6 ratios versus that assessed from WGS/OGM data. Thus, based on the WGS and OGM data, we found that the virus had become integrated in five of the six cell lines.

**Figure 3.**
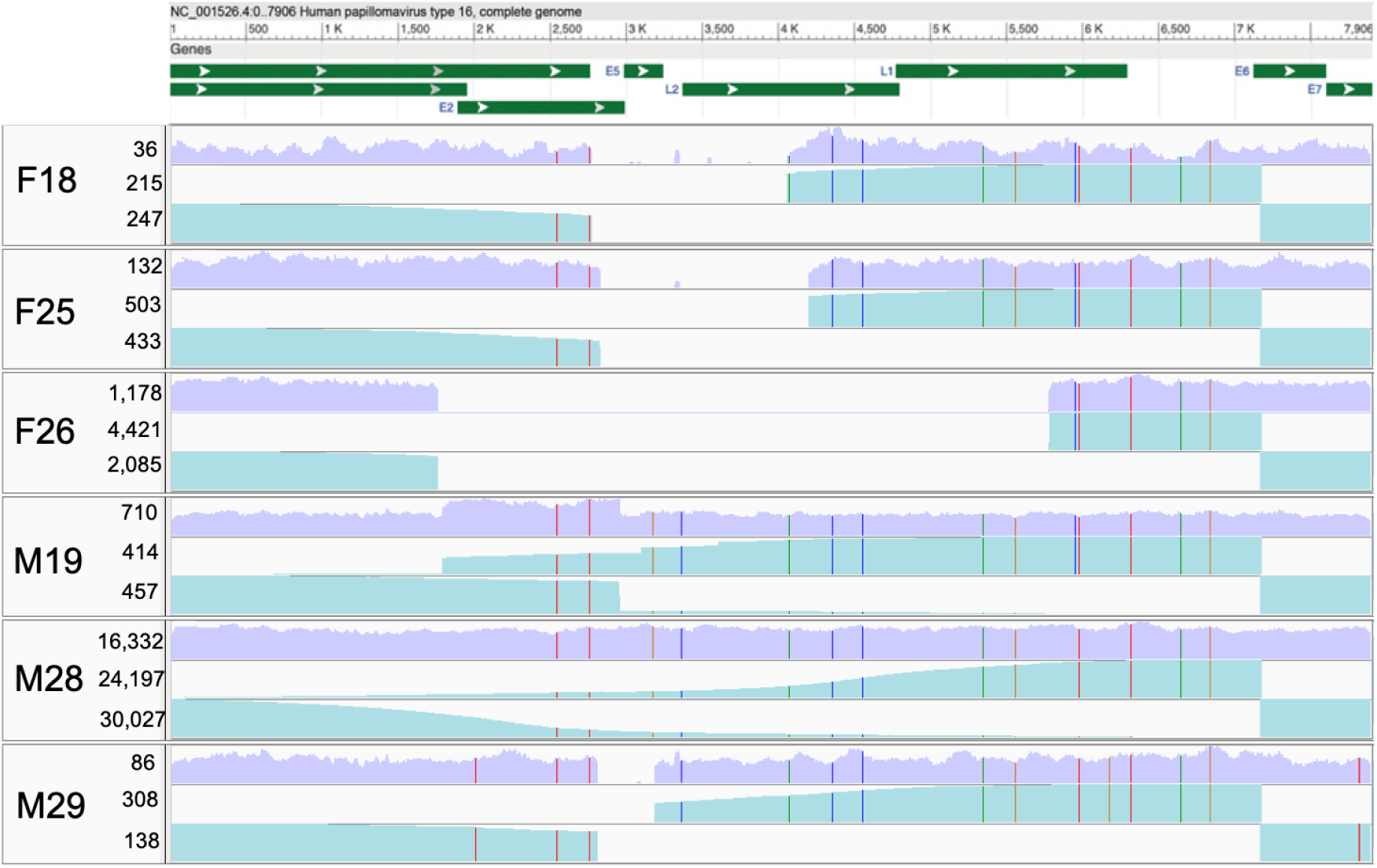
WGS and anchored PCR analysis of HPV genomes in transformed keratinocytes. Shown underneath a map of the HPV16 genome are sequence coverages as a function of genome position of virus in the six cell lines examined in this study. Coverage was determined either from the total WGS read counts at each position of the virus (violent panels) or from anchored PCR read counts (teal, from the E6 primer, second row of each panel, and from the E6v2 primer, third row in each panel). Numbers immediately to the left of each panel denotes the maximum coverage value for the adjacent sample. Positions of single nucleotide polymorphisms relative to reference HPV16 (NC_001526.4) in each virus are designated, color coded to indicate the nucleotide substitution (T, red; C, blue; A, green; G, orange).

To confirm and further characterize the status of the virus in the six cells lines, we modified the previously published LUMI-PCR technique (26) by adapting it to long-distance anchored PCR in combination with long read PacBio sequencing (see Figure S1). Specifically, we sheared DNA from the late passage cells to approximately 20 kb fragments onto which we ligated a half adapter, that is, a duplex oligonucleotide with a single strand 5’ extension whose sequence templates the subsequent PCR primer site. We then initiated two separate PCR reactions on the fragmented genomic DNA, with one primer targeting the half adapter and the other targeting the E6 coding region, using E6 primers of opposite orientation for the two separate PCR reactions. After amplification, we added hairpin oligonucleotides for initiating PacBio sequencing and subjected the samples to long range PacBio sequencing. For all cell lines, the resulting sequencing reads covered the entire residual HPV genome and, for the integrated viruses, up to ten Kb of human sequence abutting the viral integration (Figures 3, S2). Results from that analysis confirmed the HPV integration status and site of integration of the virus in all six cells lines and clarified the local structure of the virus in those lines in which it had integrated.

Consistent with previous results from tumor tissue, the site of integration of the virus into the genome in every cell line showed significant focal amplification and other structural rearrangements (Figure 4). In the simplest cases, the virus was captured at the junction of an intrachromosomal translocation that resulted in a duplication of the chromosomal segment bracketed by the translocation endpoints (F19 and M29). In a more complex case (F26) the virus integrated juxtaposed to a 90 kb fragment of chromosome 1 inserted into chromosome 12. The 200 kb region of chromosome 12 spanning the chromosome 1 and virus insertion was duplicated in tandem six times. The 11 kb of chromosome 12 immediately adjacent to the viral insertion along with the viral fragment and 12 kb of the chromosome 1 sequences on the opposite side of the viral insert excised from the chromosome as a 42 Kb human-viral hybrid episome, amplifying approximately 30-fold in the cell line. The focal amplification at the sites of integration in the other two cell lines exhibited a complexity intermediate between those two extremes.

**Figure 4.**
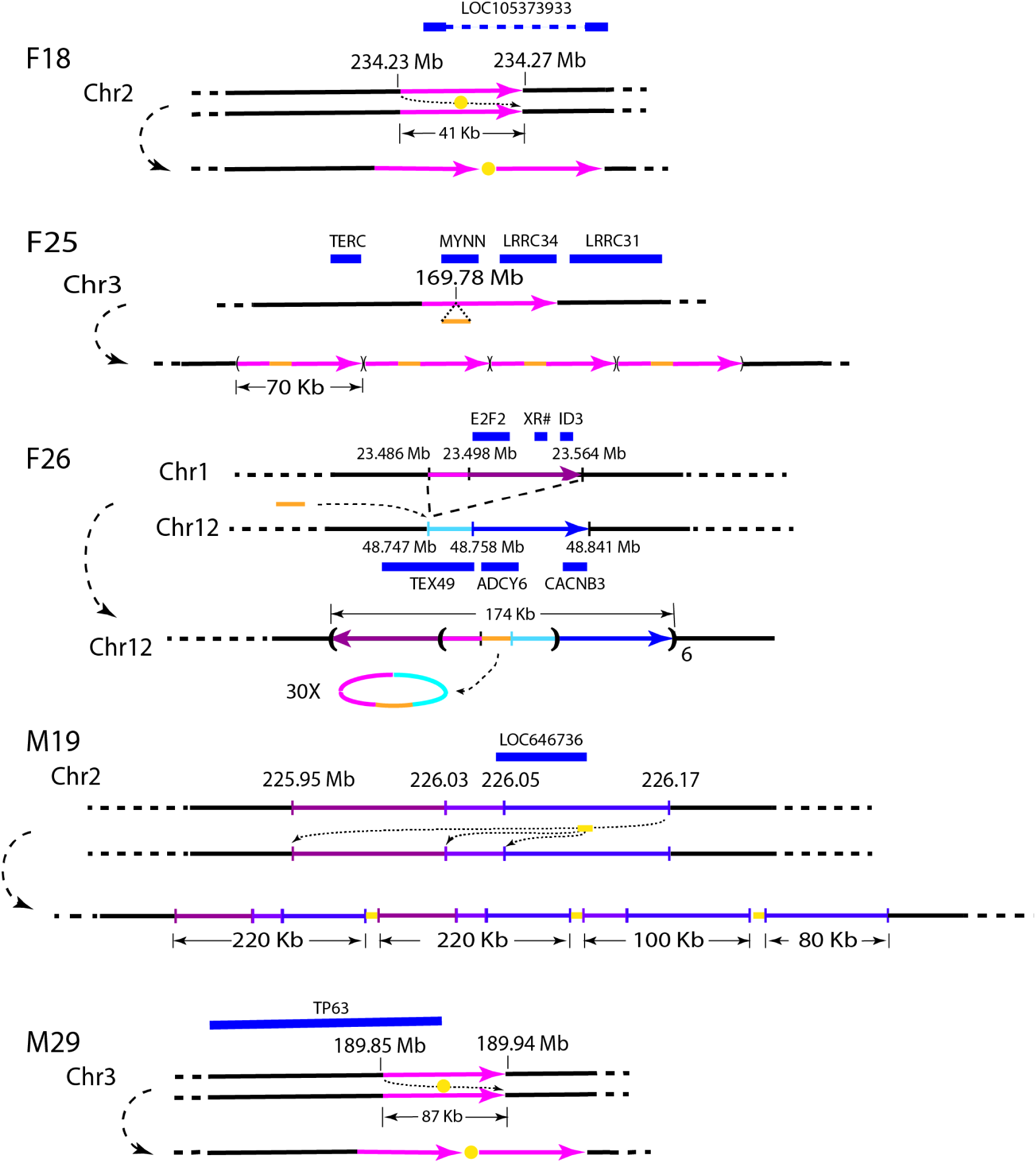
Structure of integration sites. Shown are diagrams of the regions surrounding the sites of viral integration in five cell lines. Virus is shown as a yellow bar or dot and regions of the genome that become duplicated following integration are shown in color. The upper portion of each diagram indicates the location of integration, with nearby genes shown above, while the lower portion represents the local structure following integration and focal amplification. Diagrams are not to scale.

### Substantial structural rearrangements occur genome wide prior to, but not after, viral integration

OGM performed on cells from approximately every tenth passage of the cell lines allowed us to pinpoint when virus integration occurred during the propagation of the cell line and, at the same time, assess the accumulation of structural variants over the course of propagation. Figure 5 shows a set of Circos plots from six separate passages distributed over the course of approximately sixty passages of line F18. Whole genome short read sequencing from passage 44 documented virus insertion at the junction of an intrachromosomal translocation between positions 234,232,527 and 234,273,679 on chromosome 2, an assignment confirmed by anchored PCR (Figure S2). Moreover, OGM identified an insertion/duplication at that precise position in cells from passage p44. The same exact insertion is seen in all the other passages examined except for p12. Accordingly, we conclude that the virus became stably integrated between passages 12 and 19 in this cell line.

**Figure 5.**
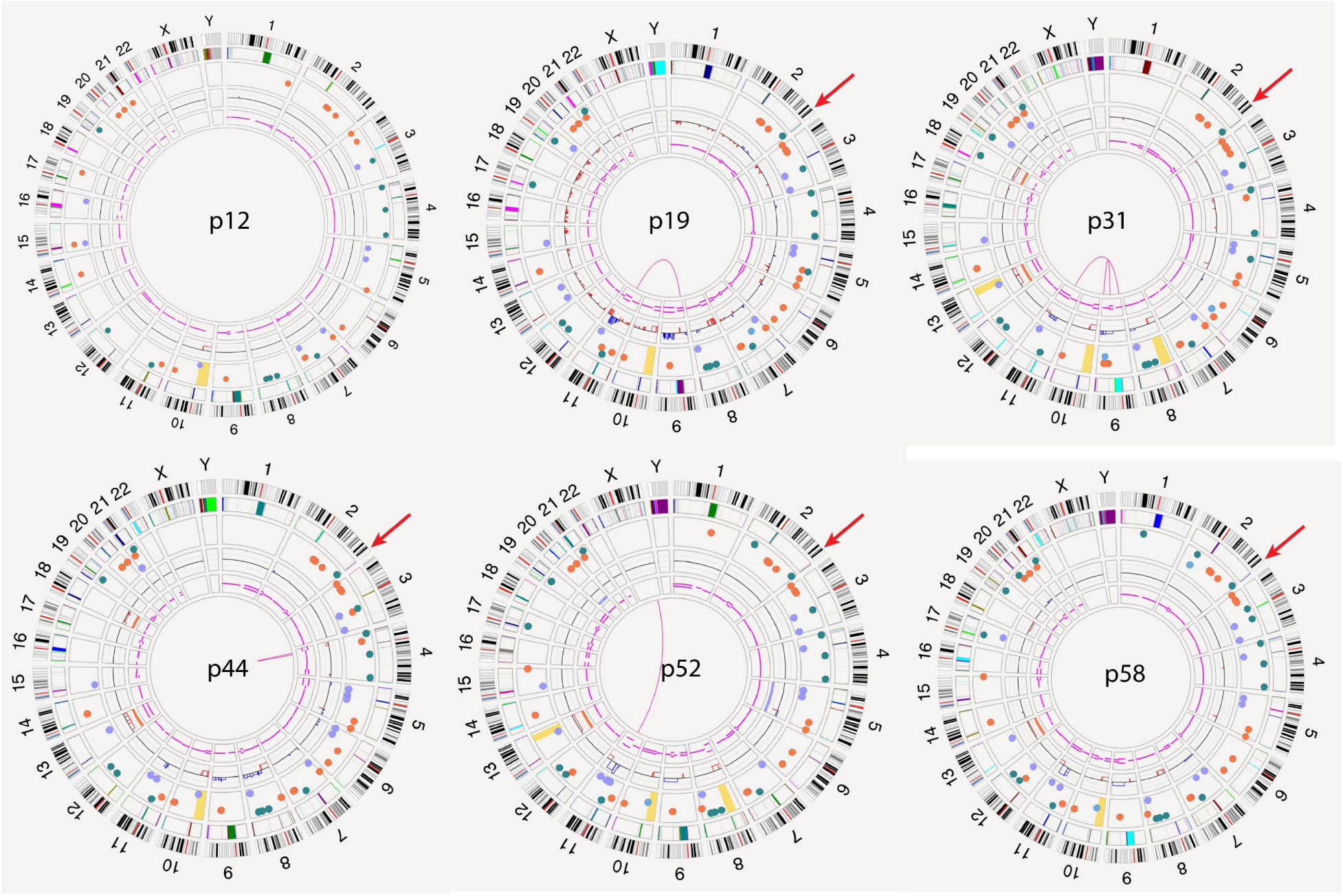
Accumulation of structural variants as a function of time in culture. Circos plots of genomes of cells from the F18 cell line taken at the indicated passage number, showing translocations and inversions in the center, copy number on the inner rings and insertions (green), deletions (orange) and duplications (light blue) on the third most outer ring. Chromosomes are ordered sequentially in the outer ring on which are indicated cytological banding patterns and the centromere (red bar). Red arrow indicates the site of virus insertion.

The circos plots also depict the accumulation of structural variants over the course of propagation of the cell line. Such variants include insertions, deletions, duplications of 500bp or greater in length as well as translocations and segmental aneuploidies of larger sizes. Because we did not have access to untransformed keratinocytes or other tissue or blood samples from the tonsil donor for each cell line, we were unable to distinguish private polymorphisms that preexisted in the donor sample from somatic structural variants that arose prior to passage 12. Numerous prior studies have documented that each individual carries 20-40 such structural variants as private polymorphisms, and we detected approximately 30 structural variants that were both present at an allele frequency of approximately 0.5 and detected in all the passages examined. Accordingly, we cannot determine the number of de novo SVs that arose prior to passage 12 and we have assumed that most of the SVs, except translocations and segmental aneuploidies, present at the earliest examined passage of each cell line are private polymorphism. However, we can calculate the number of SVs that arose in the intervals between each of the subsequent passages we examined. This is shown in Figure 6, which presents the cumulative accumulation of persistent SVs in each cell line as a function of passage number. For those cell lines in which virus integration occurred, the passage at which integration was first detected in that line is also indicated.

**Figure 6.**
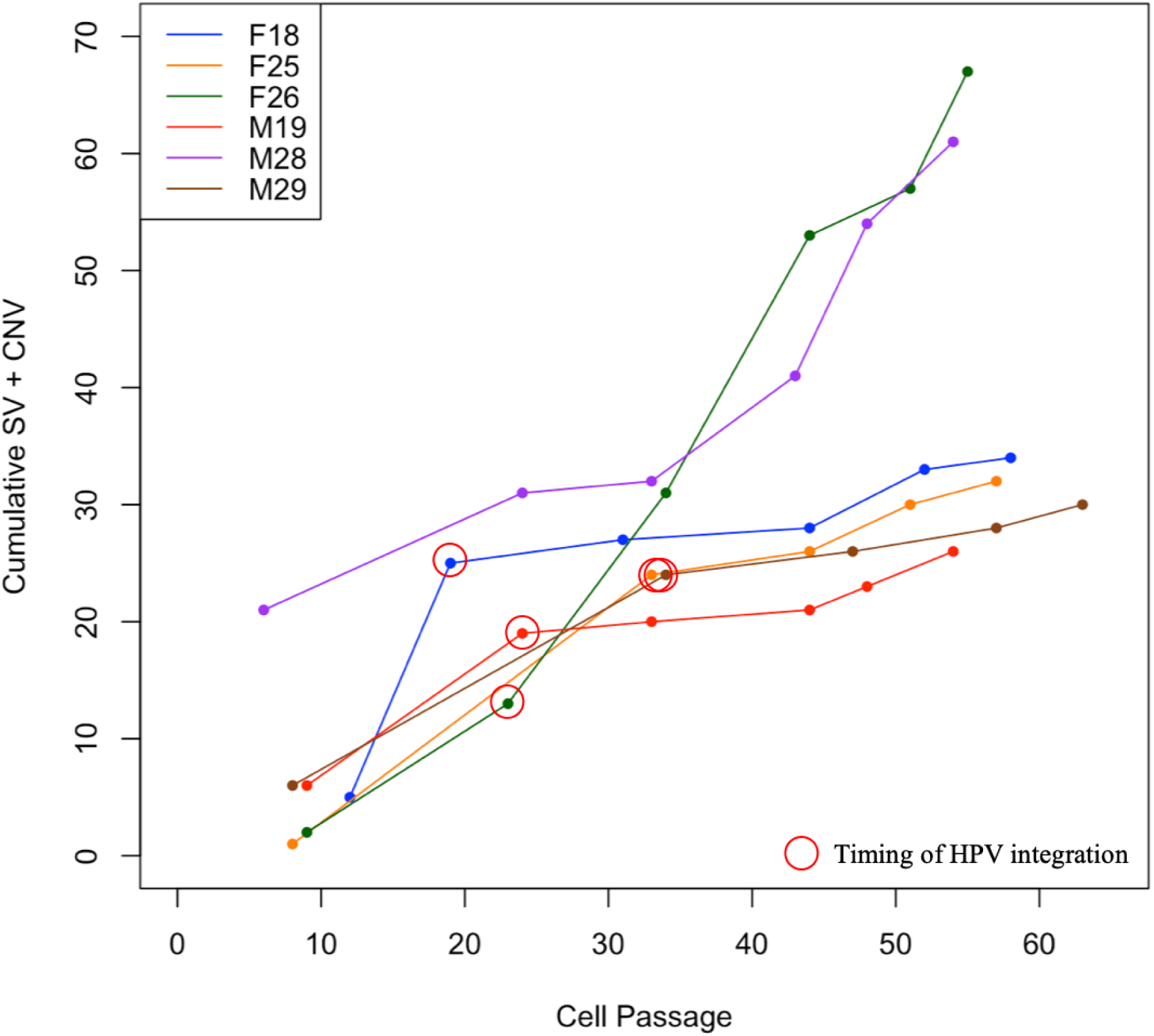
Genome Instability Precedes Viral Integration. The HPV transformed cell lines indicated were passaged and sampled at the indicated passage numbers to determine HPV integration status and the accumulation of structural variants (CVs and SVs). Red circles indicated the passage at which integration was first detected.

The data in Figure 6 suggest several characteristics regarding viral integration and SV accumulation. First, viral insertion occurred relatively early in the propagation of the culture, generally by around the 20^th^ passage. This is consistent with the data obtained from E2/E6 ratios. Second, all the lines accumulated SVs at essentially equivalent rates prior to viral insertion but for four of the five lines with insertions, the rate significantly decreased after insertion. The exceptions are the line with solely episomal HPV and the line with the vast majority of the viral genomes persisting as episomal human-virus hybrids. Accordingly, we conclude that viral integration does not promote genome instability or accumulation of SVs but rather may stabilize the genome.

### Integration is associated with more tumorigenic transformants

Our previous studies suggested that HPV+ tumors with integrated virus led to poorer outcomes than did those tumors with only episomal virus. To explore whether this correlation persists in our transformed cell lines, we performed subcutaneous injections in nude mice of later passage cells from a cell line with integrated virus (M19p44) and the one with solely episomal virus (M28p44). Prior to injection, one culture of each cell line was treated with the tobacco carcinogen dibenzo[def,p]chrysene-11,12-dihydrodiol 13,14-epoxide (DB[a,l]PDE), since prior work had indicated that such treatment enhanced the tumorigenicity of HPV transformed cell lines (31). As evident from Figure 7, both the treated and untreated M19 cells formed visible and palpable growths that expanded in volume for four weeks post injection and then decreased in size at later times. Treated cells generated slightly larger tumors but not at statistical significance. In contract, the mass from the injected M28 cells, both treated and untreated, steadily decreased in volume, essentially disappearing by the end of the observation period. These observations were consistent with our previous suggestion that HPV+ tumors with integrated virus are more aggressive than those with only episomal virus.

**Figure 7.**
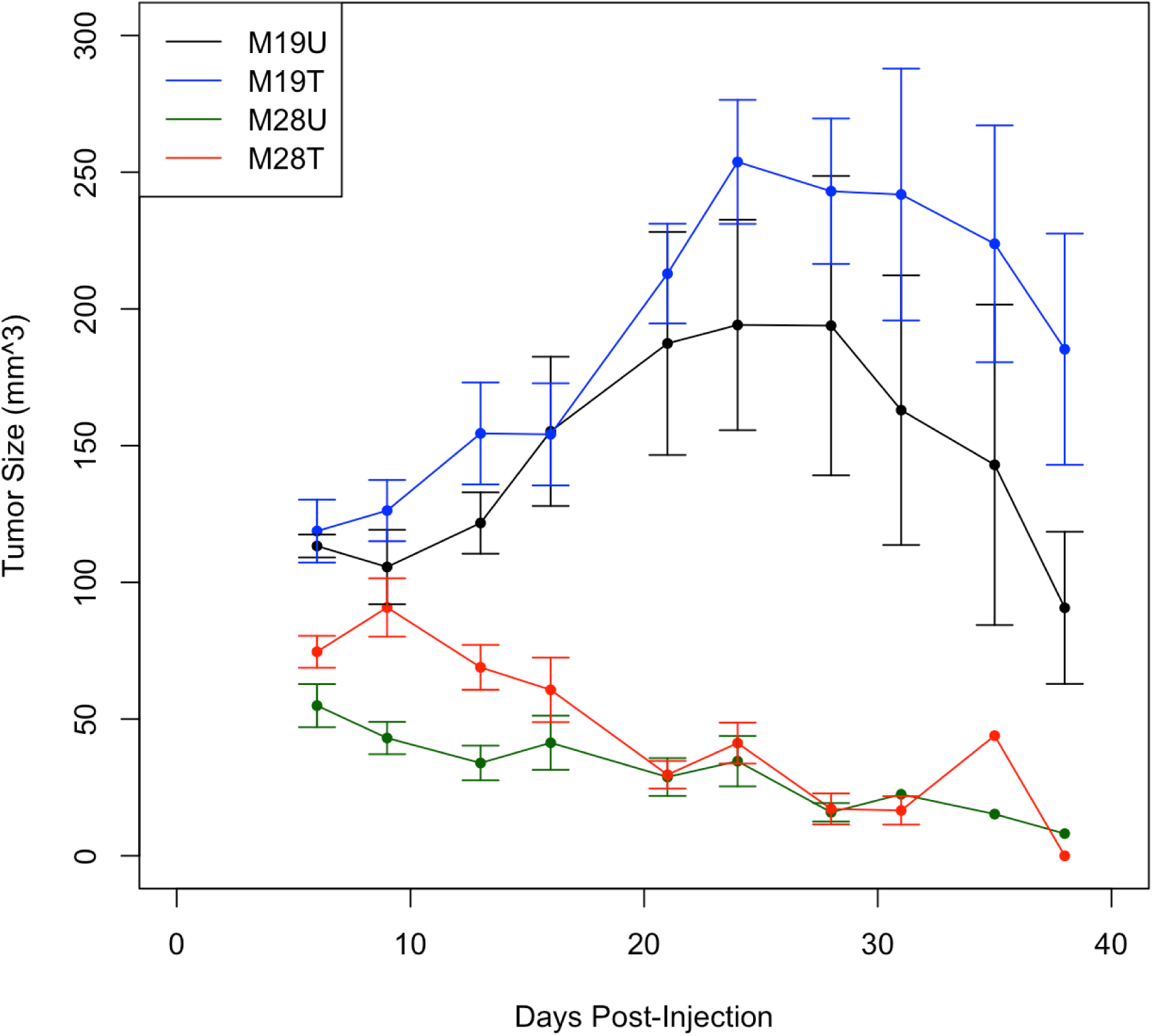
HPV Transformants with integrated virus are more tumorigenic than those with episomal virus. Cells from lines M19 (passage 44) and M28 (passage 43) were cultured for three days in E-media in the presence (T) or absence (U) of DB[a,l]PDE. Cells (10^7^) were suspended in PBS and injected subcutaneously into either flank of five nude mice for each condition. Growth was monitored by measurement of dimensions of the growth over time. Values are the means with standard deviation of the five growths from each sample.

## Discussion

### Viral integration does not cause increased genome instability

Our previous studies established a correlation between increased genome instability and viral integration in OPSCC tumors (24). This raised the question as to causality: does integration induce instability or does instability lead to integration? To address this question, we examined HPV transformed human tonsillar keratinocytes in culture and monitored accumulation of structural and copy number variants as well as viral integration during the propagation of the transformants. The results presented in this report examining the timing of genome instability and viral integration suggest that integration does not lead to instability. Rather, we found that genome instability preceded integration and diminished subsequent to integration.

The results regarding HPV integration and genome instability for HPV transformed keratinocytes in vitro exhibit both similarities with and differences from those observed for HPV+ OPSCC tumors. First, we observed focal rearrangements and amplifications at the site of viral integration in the cell lines with features quite similar to those observed at the site of HPV integration in OPSCC tumors. Second, as seen in vivo, integration appears to occur randomly, both in the viral genome and the human genome. Third, also as seen in vivo, the viral genome can persist in the cells in one or more of several different states – as an intact episomal genome, as a hybrid human/viral episome or as a fragment integrated into the host genome. Nonetheless, the major significant difference between our in vitro culture and human tumors is that we observed a continual increase in accumulation of structural variant both during the early passages and, in the lines with episomal virus, in the later passages as well. This contrasts with our previous in vivo results in which tumors with episomal virus showed little or no accumulation of structural variants.

Our observation that genome instability precedes viral integration is consistent with previous studies establishing that DNA double strand breaks in both the host and viral genome initiate integration through non-homologous end joining or microhomology directed recombination (15,32–34). Accordingly, agents that induce double strand breaks, such as oxidative stress or ionizing radiation from γ- or proton-irradiation, potentiate capture of extrachromosomal DNA (35,36). Our results suggest that initial transformation of keratinocytes by HPV and the stress of adaptation to in vitro culture conditions result in DNA damage leading to the accumulation of SVs and other somatic mutations and in some cases to the capture of a portion of the episomal viral genome. That the accumulation of structural variants does not continue after integration clearly demonstrates that viral integration does not cause DNA damage and accumulation of structural variants. Thus, these results are inconsistent with a model in which integration results in inactivation of the E2 gene with attendant increase in expression of E6, with the consequence of increased DNA damage. Why structural variants continue to increase in the line with only episomal DNA or the line with integrated and hybrid episome is unclear. One significant difference between those two lines and the integrated lines is the copy number of E6 and E7, suggesting that the exceptionally high levels of these genes may be responsible for increased DNA damage. However, this will require further evaluation.

### Viral integration is associated with increased tumorigenicity

We previously suggested that tumors with integrated virus were more aggressive than those with episomal virus. Consistent with that previous observation, we found that cells with integrated virus generated tumor masses upon subcutaneous injection into nude mice whereas the cells with only episomal virus did not. In a parallel experiment, we pretreated those same cells with the carcinogen dibenzo[def,p]chrysene-11,12-dihydrodiol 13,14-epoxide (DB[a,l]PDE) followed by subcutaneous injection in nude mice, since earlier studies using oral keratinocytes showed cooperation between HPV and tobacco carcinogens (31,37,38) and the synthetic carcinogen 4-nitroquinoline-N-oxide (4NQO) has been shown to be a potent co-factor in driving HPV associated HNSCC in HPV16 transgenic mice (39). With these pretreated cells, we also observed that the cells with integrated, but not episomal, virus induced tumors, with the pretreated cells with integrated virus showing increased in tumor volume relative to the untreated cells. We do not yet understand why cells with integrated virus appear more tumorigenic than those with only episomal virus. Certainly, integration is not the only factors distinguishing more aggressive HPV+ OPSCC from less aggressive cases: only 20% of HPV+ OPSCC exhibit poorer outcomes while 70-80% of those tumors have integrated virus. One possibility is that the structural variants arising in integrated virus can affect oncogenes or tumor suppressor genes well removed from the site of integration, as we observed in our previous study. Nonetheless, our the results presented here reinforce the notion that viral integration, or lack thereof, may prove to be a useful prognostic indicator for cases of HPV+ OPSCC.

## Acknowledgements

The authors would like to thank Dr. Craig Praul in the Genomics Core Facility of the Penn State Huck Institute for performing long range sequencing on the PacBio Sequel IIe System.

## Data Access

Bionano variant calls and mapped reads for our samples can be downloaded from https://www.datacommons.psu.edu/commonswizard/MetadataDisplay.aspx?Dataset=6286.

## Author Contributions

CM, DG and JRB conceived the experiments and supervised their execution. KC, CT, EM and DG performed the genomic studies. SA and SC performed cell culturing and histochemistry. LS, KS, KC and JRB analyzed the genomic data. KC and JRB wrote the manuscript, which was edited by DG and CM.

